# Anthelmintic resistance against benzimidazoles and macrocyclic lactones in strongyle populations on cattle farms in northern Germany

**DOI:** 10.1101/2024.12.20.629641

**Authors:** Paula Ehnert, Jürgen Krücken, Stefan Fiedler, Fabian Horn, Christina S. Helm, Ann Neubert, Wiebke Weiher, Werner Terhalle, Stephan Steuber, Ricarda Daher, Georg von Samson-Himmelstjerna

## Abstract

Anthelmintic resistance (AR) in cattle gastrointestinal nematodes (GIN) is an increasing global concern, with low to moderate levels recently documented in Central Europe. This study reports on resistance against both macrocyclic lactones (MLs) and benzimidazoles (BZs), highlighting that AR is spreading. The Fecal Egg Count Reduction Test (FECRT) remains the primary tool for AR assessment, yet differing methodologies and recent guideline updates complicate resistance interpretation across studies. Statistical methods, such as Bayesian approaches used by eggCounts and bayescount, yield varying confidence intervals, further influencing results. Notably, the nemabiome analysis identified *Ostertagia ostertagi* and *Cooperia oncophora* as predominant species in the region, though unexpected diversity among farms with additional GIN species occurring sometimes even at high frequency, suggests morphological analysis of coprocultures may underestimate species prevalence. Detecting AR against both drug classes on some farms underscores the urgency of implementing sustainable strategies, such as targeted selective treatment and combinations of anthelmintics with different mode of action, to prevent scenarios of multi-drug resistance observed elsewhere. Effective resistance management requires immediate discussions with veterinarians and stakeholders to steer toward informed, preventive measures in cattle farming.

## Introduction

For grazing livestock, infections with helminths are inevitable^1^ and may lead to considerable animal welfare problems^2^. Associated economic losses attributed to production losses (∼80%) and treatment costs (∼20%) in 16 European countries plus Israel and Tunisia were estimated to be 1.8 billion € for ruminant livestock and about1.4 billion for cattle alone^3^. Production losses due to gastrointestinal nematodes (GIN) were assumed to be as high as 686 million €. In the absence of vaccines and highly effective control measures based on farm management that do not result in high workloads, commercial farming widely relies on anthelmintic treatments to control strongyle infections^4^.

Cattle are particularly susceptible to infection in their first and second grazing seasons while older animals develop a partial immunity and often show low egg shedding associated with subclinical symptoms only^5^. Young stock is not only the age group that suffers most from infections with GIN^6^ but is also best suited to evaluate the efficacy of control measures including treatment with anthelmintics due to their high egg shedding intensity^7^. Although, it is assumed that a combination of substances that act against the same helminths but have different mechanisms of action may delay the development of resistance to the classes of substances included, there are some concerns by the approving authority, the European Medicines Agency (EMA), that the practice of combination could lead to development of simultaneous resistance to multiple anthelmintic classes^8^. However, in contrast to several other regions in the world with industrial cattle farming, combinations of anthelmintics with the same-spectrum of efficacy to use synergistic efficacies^9^ are not licensed and available yet in the EU. Therefore, farmers mostly have to rely on the efficacy of mono-drug treatments for the control of GIN infections. In South America, the USA, Australia and New Zealand anthelmintic resistance in cattle has already been identified as a major problem^9, 10, 11^. Fortunately, reports of anthelmintic resistance (AR) in cattle in Central Europe are still scare and do so far not indicate a severe problem: Previous studies from Germany^12, 13^ and Central Europe^14, 15, 16^, detected low to moderate levels of resistance against macrocyclic lactones (MLs) while resistance against benzimidazoles (BZs) was rare and has so far not been described for Germany. However, the number of studies that were conducted to monitor resistance levels is small and generally not sufficient to provide a representative picture about the current status^17^. In Sweden and Germany, no albendazole resistance in GIN was detected when tested on two and ten farms, respectively^13^. In contrast, ivermectin (IVM) resistant GIN were frequently detected in the same study on farms in Belgium, Sweden and Germany. While *Cooperia (C.) oncophora* was associated with poor efficacy in all three countries, *Ostertagia (O.) ostertagi* appeared to be still fully susceptible in Belgium and Germany but not in Sweden. For cattle farms in northern Germany, BZ susceptibility and frequent occurrence of ML resistance was also confirmed using the egg hatch and the larval migration inhibition tests, respectively^18^. In Belgium and Germany, El-Abdellati, et al.^19^ found reduced efficacy of MLs on 39% of 84 farms with *C. oncophora* identified as the species involved in resistance according to coprocultures post-treatment. Geurden, et al.^14^ confirmed AR in GIN on 40 farms from the countries Germany (1/12 farms), UK (3/10), Italy (10) and France (3/8) employing moxidectin (MOX) and IVM. Similarly, Charlier, et al.^3^ reported IVM resistance in GIN on some farms in France but not Italy while GIN on all tested farms were susceptible to BZs. In contrast, a study from Ireland also detected frequent resistance against fenbendazole (FBZ), levamisole, IVM and MOX^20^.

Based on larval cultures, the most frequently reported cattle GIN are *C. oncophora* and *O. ostertagi*^13^, although identification of larvae to the species level is difficult and *Cooperia* spp. and *Ostertagia* spp. would probably be more correct. More diverse species communities were reported by others and these included multiple *Cooperia* spp., *Trichostrongylus* spp., *Oesophagostomum* spp., *Bunostomum phelebotomum* and *Haemonchus contortus*^21^. In the last years, deep amplicon sequencing based approaches targeting the internal transcribed spacer 2 (ITS-2) region of the ribosomal RNA gene locus (nemabiome)^22^ have been used to characterize strongyle communities in cattle^23^, bisons^24^, horses^25^ and sheep^26^.

The aim of the present study was to evaluate the efficacy of MLs with a focus on eprinomectin (EPR) and the BZ FBZ on GIN communities using the fecal egg count reduction test (FECRT) on cattle farms in north-east Germany. Data were evaluated using different statistical approaches to calculate 95% confidence intervals (CIs) and interpretations were based on both the original guideline for the FECRT^27^ and the recently revized guideline^28^. Furthermore, the project aimed to characterize the GIN communities before and after treatment using nemabiome analysis.

## Materials and methods

### Ethics statement

The present study did not qualify as an animal experiment requiring ethical approval according to the responsible authority Landesamt für Arbeitsschutz, Verbraucherschutz und Gesundheit (LAVG) Brandenburg (AZ-7221.3-14163_21).

### Fecal egg count reduction test and sample collection in cattle farms

The study was conducted in cattle farms located in Brandenburg, Mecklenburg-Western Pomerania, and Saxony-Anhalt between 24.08.-14.12.2021. Following discussions with local animal welfare authorities, it was confirmed that the research on cattle farms (as decided for a previous study on sheep by Krücken et al.^29^ did not qualify as an animal experiment requiring ethical approval according to the responsible authority Landesamt für Arbeitsschutz, Verbraucherschutz und Gesundheit (LAVG) Brandenburg (AZ-7221.3-14163_21).

Farms were contacted using recommendations from local veterinarians. Additionally, a call for study participation was shared through Biopark e.V., the East German edition of Bauernzeitung, and the RBB Cattle Breeding Association Berlin-Brandenburg.

The cattle farms selected for this study fulfilled the COMBAR guidelines^30^ following published suggestions^7^ and were in agreement with the criteria published in a recent guideline^28^. Each farm had to have between 10 and 40 first- and second-year grazing cattle that had not been dewormed for at least 6 weeks prior to the study. Additionally, facilities were required to capture and immobilize the animals twice during the study.

Since all except one of the farms had at least 40 animals that could be included, the cattle were divided into two groups of 20 animals per farm. One group was treated with a subcutaneous EPR and the other with an oral FBZ formulation. On farm 5, only one group treated with EPR was included since the number of available animals was too low to test efficacy of both drugs. The dosages were calculated according to the animal weight, which was determined using either a livestock scale or by measuring the chest circumference with an Animeter®. Specifically, the subcutaneous injection used Eprecis® (20 mg/ml) (CEVA Tiergesundheit GmbH) containing EPR at a dose of 0.2 mg/kg body weight, while the oral Panacur® Suspension 10% (Intervet Deutschland GmbH) with FBZ was administered at a dose of 7.5 mg/kg body weight. As requested by the farmer of farm 2, instead of the EPR treatment the IVM preparation Chanectin® 0.5% solution (Alfavet Tierarzneimittel GmbH) was administered as a pour-on application at a dose of 500 µg/kg body weight. Those last mentioned topically treated animals were kept separate from untreated or differently treated animals for 14 days after treatment until the control sampling. Individual fecal samples were collected from the rectal ampulla on the day of treatment and again 14 days later.

### FLOTAC analysis

Fecal samples were analyzed using the FLOTAC system^31^ according to the manufacturer’s instructions. Individual fecal samples were weighed (10 g), suspended in 90 ml of tap water, filtered through a metal sieve and 11 ml of the suspension were transferred into a centrifugation tube. Samples were centrifuged at 170×g for 2 minutes and the supernatant discarded. The sediment was resuspended in 11 ml saturated NaCl solution (specific density 1.2), then loaded into the FLOTAC device, which was centrifuged at 120×g for 5 minutes. Both chambers of the FLOTAC reading disk were analyzed under a microscope, with each observed egg representing one egg per gram (EPG).

### Farmer survey

In addition to the FECRT, a paper-based questionnaire was completed by farmers, collecting demographic data (e.g. herd size, age composition, cattle breeds and usage), management practices (housing, purchases, quarantine) and information about anthelmintic use (product name, frequency, application methods) as well as experiences with fecal examination and alternatives to chemical deworming, and finally findings of parasites at slaughter. For more details see Krücken et al.^29^.

### Larval cultures from pooled fecal samples

Fecal samples from each farm collected on the day of treatment were pooled, cultured and used to obtain third stage larvae (L3). After FECRT diagnostics, the remaining fecal material was homogenized and mixed with wood shavings. This mixture was loosely packed into glass jars, incubated at 25 °C and 80% humidity for 10 days. Larvae were harvested by filling the jars completely with tap water, inverting them into petri dishes, and allowing the larvae to migrate overnight into the area of the dishes not covered by the jars. The larvae were then transferred into centrifugation tubes and triplicate 10 µl aliquots were microscopically counted to obtain data on larval numbers. Aliquots containing approximately 1,000 L3 were stored at - 20°C.

### DNA extraction and deep amplicon sequencing analyses

DNA was extracted from pooled larval samples using the Macherey-Nagel^®^ SOIL Kit. The kit was used before to obtain DNA with low concentrations of PCR inhibitors from fecal samples to amplify nematode amplicons from eggs and larvae^32, 33^. The ITS-2 region was then amplified using specific PCR primers for the ITS-2 region NC1/NC2^34^ modified by addition of Illumina^®^ adapters in a first and reamplification with primers containing a DNA barcode for bioinformatic assignment of sequences to specific samples (farms and sampling time point) in a second, limited-cycle PCR^23^. In order to avoid problems with calculation of correction factors, which is not possible if the same base is detected for all positions on the Illumina Flow Cell during primer sequencing, zero to three random bases (N0-3) were inserted between the NC1/NC2 sequences and the Illumina adapters, as detailed by Avramenko, et al.^23^ (Table S1). For details regarding amplification, reamplification, purification and concentration determination of PCR products in the different steps see Krücken, et al.^29^. The ITS-2 PCR products were sequenced on an Illumina® MiSeq with V3 chemistry (2×300 bp) aiming to obtain at least 10,000 reads per sample.

The sequencing data were assessed for quality control using FastQC software and analyzed using the Nemabiome workflow based on the dada2 pipeline^35^ and the nematode ITS-2 database version 1.3^36^. The final amplicon sequence variant (ASV) table provided an overview of the detected sequence variants, which were assigned to specific nematode species. Then, correction factors for host species-specific nematode populations were applied^23^. Analysis was performed in parallel with analysis of sheep samples as recently detailed in Krücken, et al.^29^.

### Statistical analyses

For all datasets, statistical analysis to calculate the FECR was performed using the eggCounts package to estimate the FECR and the 90% and 95% CrI. Using the bayescount package, it was only possible to calculate the 90% CrI and no estimate of the FECR. Results of the eggCounts package were interpreted based on the original guideline for the FECRT^27^ and the revised guideline^28^. The bayescount results were only interpreted using the revised guideline since the method doess not provide an estimate for the FECR. The interpretation of the data according to the two guidelines is summarized in Table 1 and has been illustrated in Fig. 1 in Denwood, et al.^37^.

**Fig. 1.**
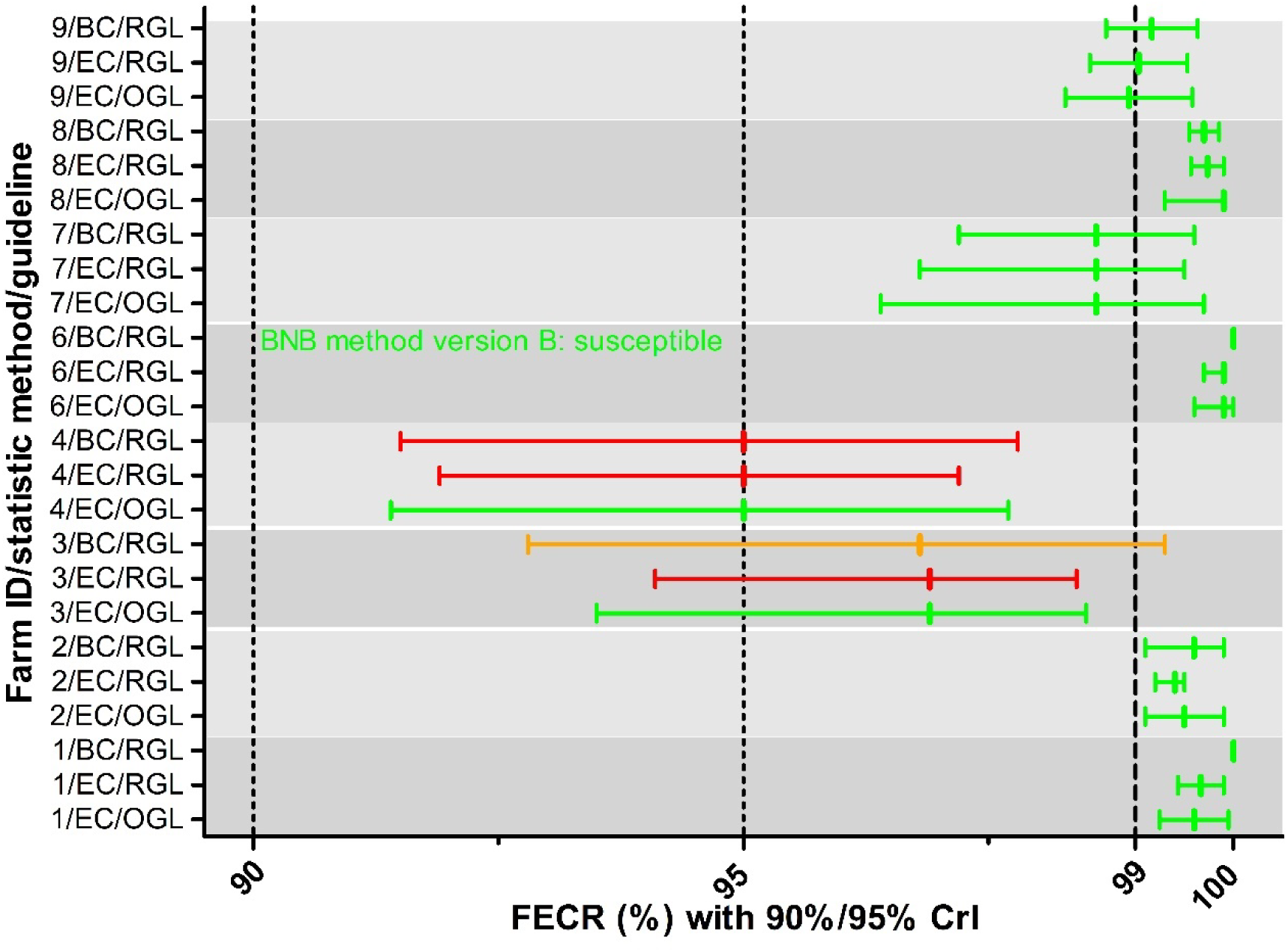
Fecal egg count reductions (FECRs) with credible intervals (CrIs) for fenbendazole. The CrIs were calculated using either eggCounts (EC) or bayescount (BC). Interpretation of the results was based on the revised guideline (RGL) for the FECR test^28^ with 90% CrIs shown, or on the original guideline (OGL)^27^ with 95% CrIs shown. Since bayescount does not calculate an estimate of the FECR, the value was calculated from mean egg counts pre- and post-treatment. Colors indicate the status assigned to the strongyle communities: green, susceptible; red, resistance; light red, beginning resistance; orange, inconclusive / suspected resistance. For one farm, with an EPG of zero for all animals post treatment, bayescount was not able to calculate a 90% CrI. In this case the BNB method version B was used to assign a status to the community^37, 40^.

**Table 1.** Comparison of criteria for interpretation of data from the fecal egg count reduction (FECR) test. CL, confidence limit; n.r., not relevant; beginning resistance is a subcategory within the category resistance.

| Original guideline (Coles et al.)<sup>27</sup> |  |  |  |
| --- | --- | --- | --- |
| Interpretation | Lower 95% FECR CL<sup>a</sup> | FECR estimate | Upper 95% FECR CL<sup>a</sup> |
| Normal | >90% | >95% | n.r. |
| Suspected resistance | <90% | >95% | n.r. |
| Suspected resistance | >90% | <95% | n.r. |
| Resistance | <90% | <95% | n.r. |
| Revised guideline (Kaplan et al.)<sup>28</sup> |  |  |  |
| Interpretation | Lower 90% CL<sup>a</sup> | FECR estimate | Upper 90% FECR CL<sup>a</sup> |
| Susceptible | >95% | n.r. | >99% |
| Inconclusive | <95% | n.r. | >99% |
| Beginning resistance | >95% | n.r. | <99% |
| Resistance | <95% | n.r. | <99% |

The FECR, along with the 90% and 95% credible intervals (CrIs), were calculated using the eggCounts package (version 2.3-2) in R 4.1.3 using a Bayesian analysis based on Markov Chain Monte Carlo analysis and raw egg count data^38^. The algorithm was applied to paired data, with consistent efficacy across the farm and no zero-inflation and an estimate for the FECR (mode of the posterior distribution) and the 95% CrI (2.5% and 97.5% quantiles of the posterior distribution) were obtained. The 90% CrI was extracted using the coda package (version 0.19–4) and results were interpreted following WAAVP guidelines^28^. In addition, the bayescount package version 1.1.0^39^ as implemented on the web portal fecrt.com (last viewed 07. October 2024) was used to calculate 90% CrIs using a second, also Bayesian statistical approach. The interpretation of the FECRT results were compared between eggCounts and bayescount statistics but for eggCounts also between the criteria suggested by the original^27^ and the revised^28^ guideline for the FECRT. While the original guideline relies on the estimate of the FECR and the lower 95% credible limit (CrL), the revised guideline completely focuses on the lower and upper 90% CrLs^27, 28, 37^. In some cases (e.g. when all post treatment EPG values are 0, bayescount is unable to calculate 90% CrI for the FECR. In these cases, the software chooses automatically the best option among various variants of Beta Negative Binomial (BNB) methods^37,40^ .

In order to quantitatively compare agreement between statistical methods and between original and revised guideline, Cohen’s κ coefficients were calculated using the CohenKappa() function from the DeskTools package 0.99.48. Interpretation of Cohen’s κ coefficients followed the suggestions introduced by Landis and Koch^41^.

The 95% confidence intervals (CIs) for frequencies were calculated as Wilson score intervals using the binom.wilson() function from epitools package 0.5-10.1. The tab2by2() function from epitools was applied to identify significant differences between frequencies using the mid-p exact test. Egg counts were compared before and after treatment using the Wilcoxon matched-pairs signed rank test in GraphPad Prism 5.03 (GraphPad Software, Inc., Boston, MA). Correction of p values for multiple testing was conducted using the p.adjust() function in basic R.

## Results

### Farms included in the study and parasite management

The majority of the included farms were from the federal state of Brandenburg (n = 7) but there was one farm included from Saxony-Anhalt and one from Mecklenburg– Western Pomerania. All farms used an extensive farm management with pasture access for the animals. Grazing land ranged from 7 to 1800 hectares. Seven of the nine farms grazed their cattle year-round, while two housed them during the winter. The average herd size was 932 animals per farm, ranging from 52 to 3521. All farms were operated commercially, primarily focusing on cow-calf production (6 farms), with fewer engaged in only beef (3 farms) and dairy production (2 farms). Farm 7 did not provide usable data. One farm (farm 1) also kept 400 sheep and 12 goats, while another (farm 8) had 70 horses. Common cattle breeds included Uckermärker (3 farms), Angus (2 farms), Holstein-Friesian (2 farms), water buffalo (1 farm) and Charolais (1 farm).

Seven out of nine farms reported deworming their animals regularly, treating the entire herd at once, while two farms dewormed on a need-only basis. Deworming was carried out by the farmer on four farms, by a veterinarian on four farms, and by both on one farm. Ivermectin was the most commonly used anthelmintic (8 farms), followed by doramectin (2 farms), EPR (1 farm), and FBZ (1 farm). Two farms determined the animal’s weight before dosing, while the rest relied on estimates. Most farms (8/9) used pour-on applications for deworming.

Only one farm had submitted fecal samples for testing prior to the study though rather to confirm the suspicion of a liver fluke (*Fasciola hepatica*) infection than GIN. The majority (8/9) considered testing useful, potentially suggesting financial or labor constraints as a limiting factor.

Regarding herd management, three farms separated newly purchased animals from the main herd, while five dewormed them before allowing access to pasture. None of them controlled the effect of quarantine or treatment before allowing pasture access. Eight of the nine farms felt well-advised by their veterinarians and well-informed about deworming.

### Fecal egg count reduction test

Fenbendazole and EPR were used on eight of the nine farms. On farm 2, IVM (pour on) was administered instead of EPR, and on farm 5, only EPR but not FBZ was tested due to the small herd size. After treatment with any drug, FECs were significantly reduced for strongyles (Fig. S1) on all farms and on farm 6 also for *Nematodirus* spp. (Fig. S2).

Reduced efficacy in strongyles against FBZ was observed on two of the eight farms (farms 3 and 4) (Fig. 1). Using eggCounts and the revised guideline, strongyles on both farms were assigned a resistant status. When bayescount was used with the revised guideline, only data for farm 4 were considered to correspond to resistance while data for farm 3 were considered inconclusive. When the original guideline was applied, the strongyle communities from both farms were considered susceptible. For all other six farms, all three methods assigned a FBZ susceptible status to the strongyle communities (Fig. 1).

For the MLs, strongyle communities on only three farms were considered susceptible to EPR with all three methods (farms 6, 7, and 9) (Fig. 2). For strongyles on farm 3, all three statistical methods indicated EPR resistance and for farm 2 all three methods revealed IVM resistance. For worm communities on the remaining four farms on which EPR efficacy was evaluated, results differed between the statistical approaches (Fig. 2). On farm 1, CrIs were very similar but interpretation was considerably different with beginning resistance for eggCounts and the revised guideline while the other two methods indicated susceptibility. On farm 4, eggCounts with the original guideline concluded that the community was susceptible, eggCounts with the revised guideline concluded resistance and bayescount with the revised guideline let to inconclusive results. However, the upper 90% CrL was exactly 99%, any further reduction would have led to the conclusion resistance for this community. For farm 5, eggCounts with the revised guideline revealed that the community exhibited beginning resistance while the other two methods suggested that it was susceptible. Finally, on farm 8, eggCounts with the original guideline assigned a susceptible status to the community whereas the other two methods considered the community as showing (beginning) resistance (Fig. 2).

**Fig. 2.**
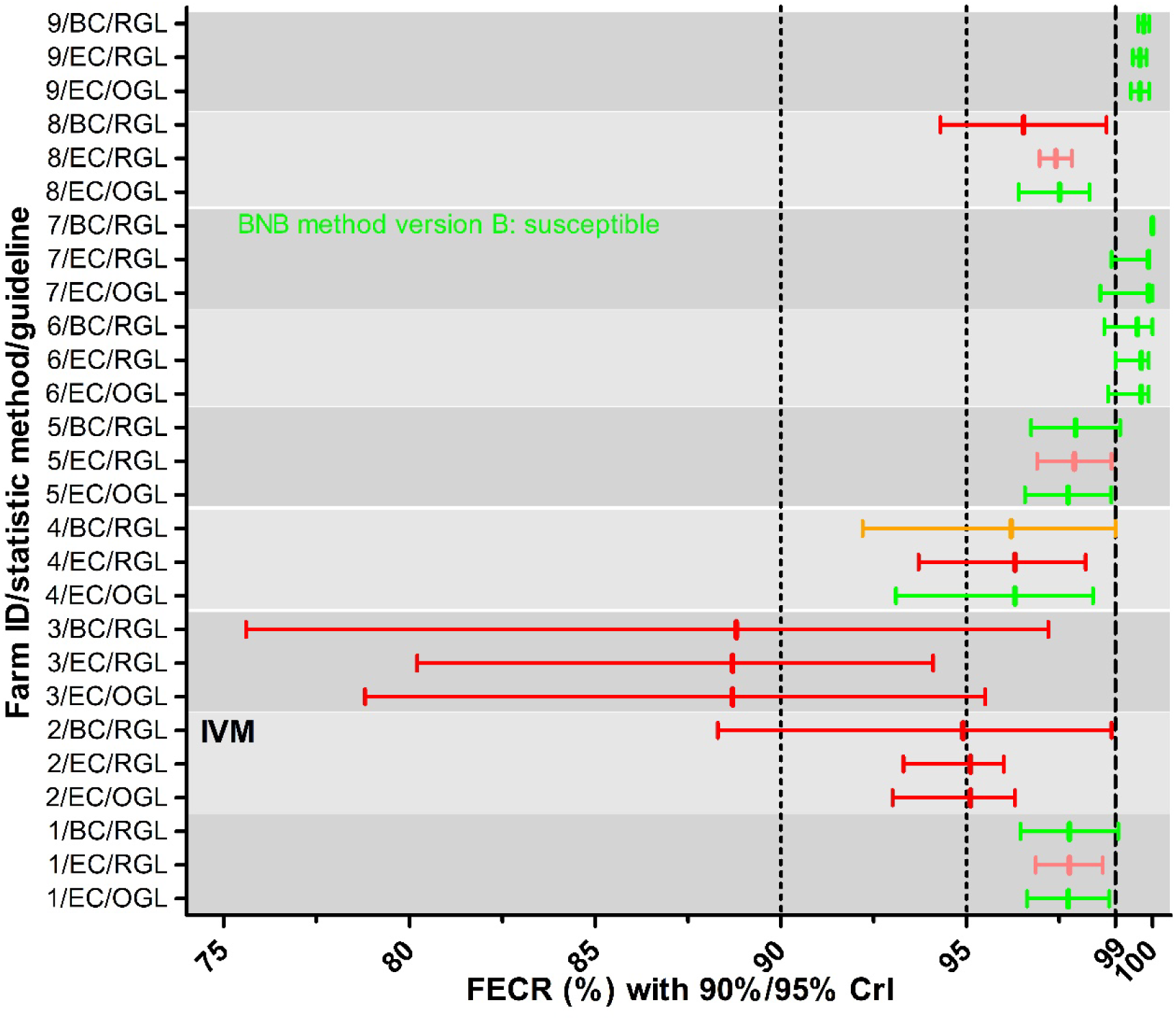
Fecal egg count reductions (FECR) with credible intervals (CrIs) for macrocyclic lactones. On all farms EPR was used to conduct the FECR test except of farm 2 where ivermectin (IVM) was used. The CrIs were calculated using either eggCounts (EC) or bayescount (BC). Since bayescount does not calculate an estimate of the FECR, the value was calculated from mean egg counts pre- and post-treatment. Interpretation of the results was based on the revised guideline (RGL) for the FECR test^28^ with 90% CrIs shown, or on the original guideline (OGL)^27^ with 95% CrIs shown. Colors indicate the status assigned to the strongyle communities: green susceptible; red, resistance; light red, beginning resistance; orange, inconclusive/suspected resistance. For one farm, with an EPG of zero for all animals post treatment, bayescount was not able to calculate a 90% CrI. In this case the BNB method version B was used to assign a status to the community^37, 40^.

On farm 6, the *Nematodirus* spp. egg count was high enough to calculate a FECR for this genus (FBZ: 392, EPR: 339 raw egg counts before treatment). The *Nematodirus* community on this farm showed resistance to FBZ and susceptibility to EPR. Interpretation of all results was identical for all three statistical methods (Fig. 3).

**Fig. 3.**
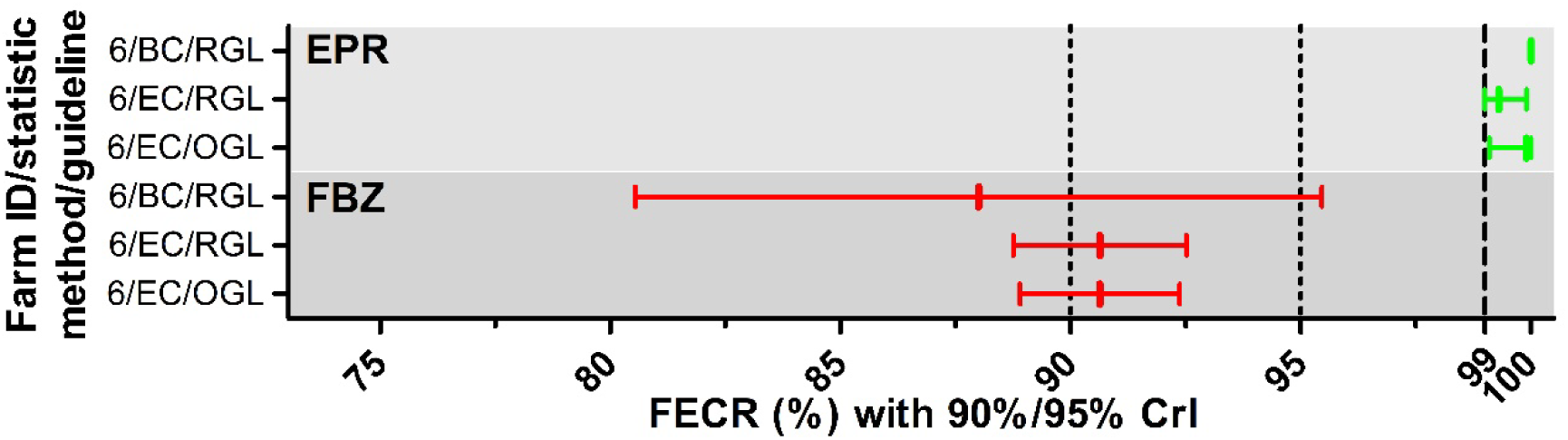
Fecal egg count reductions (FECR) with credible intervals (CrIs) for *Nematodirus* spp. The FECR test was conducted with fenbendazole (FBZ) and eprinomectin (EPR). The CrIs were calculated using either eggCounts (EC) or bayescount (BC). Since bayescount does not calculate an estimate of the FECR, the value was calculated from mean egg counts pre- and post-treatment. Interpretation of the results was based on the revised guideline (RGL) for the FECR test^28^ with 90% CrIs shown, or on the original guideline (OGL)^27^ with 95% CrIs shown. Colors indicate the status assigned to the strongyle communities: green susceptible; red, resistance.

For the strongyle nematodes including *Nematodirus* spp., no significant difference in FECR as estimated using eggCounts was observed between FBZ and EPR (Fig. 4) using the Mann-Whitney U test.

**Fig. 4.**
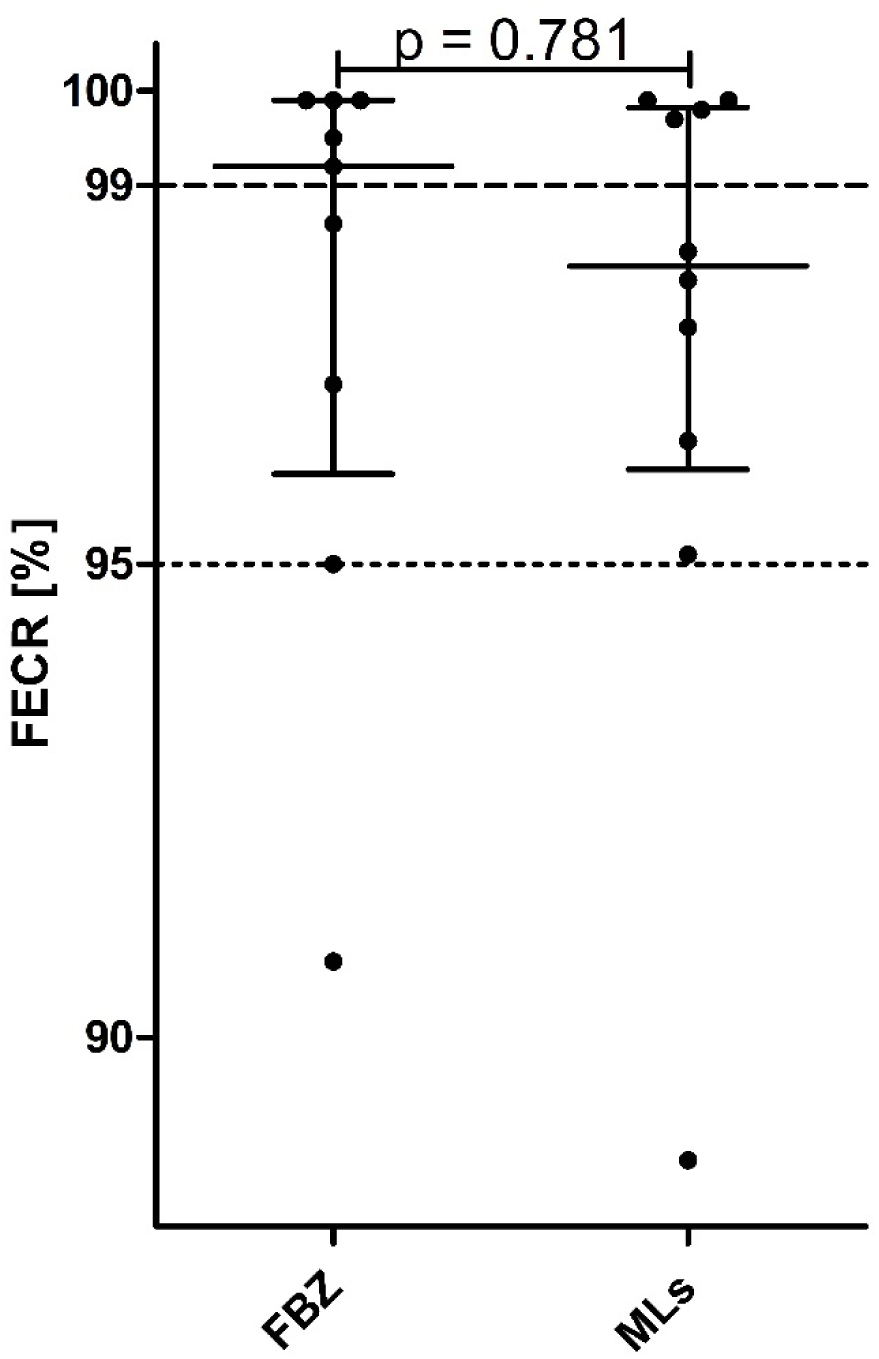
Comparison of fecal egg count reduction estimates calculated using eggCounts between fenbendazole (FBZ) and eprinomectin (EPR). Data points correspond to FECR for individual farms. Means with SD are shown. The Mann-Whitney U test was used to compare both groups.

### Comparison interpretation of the fecal egg count reduction test between statistical methods and guidelines

Using three categories for each method based on the revised guideline and treating results with the assignment beginning resistance as resistant, both methods agreed for 17/21 datasets (Table 2). Four datasets that were considered resistant by eggCounts were considered inconclusive (n = 2) and susceptible (n = 2) by bayescount. This resulted in a Cohen’s κ value of 0.518 corresponding to a moderate agreement. Comparison between eggCounts and bayescounts revealed that the 90% CrI obtained by bayescounts were significantly wider than for eggCounts (Fig. S3).

**Table 2.** Inter-rater agreement between eggCounts and bayescount results based on the revised guideline for the fecal egg count reduction test.

|  | eggCounts |  |  |
| --- | --- | --- | --- |
| bayescount | Susceptible | Inconclusive | Resistance |
| Susceptible | 10 | 0 | 2 |
| Inconclusive | 0 | 2 | 2 |
| Resistance | 0 | 0 | 5 |
| Cohen's $\kappa$ 0.518 | | | |

Comparing interpretation of the results of the FECRT between the original^27^ and the revised guideline^28^ was conducted only based on eggCounts data since bayescount does not calculate an estimate for the FECR, which is required for interpretation of results using the original guideline. Interpretation of results using both guidelines agreed for 12/21 datasets (Table 3). However, there were six datasets that were considered resistant according to the revised but susceptible according to the original guideline (Table 3). This resulted in an only fair agreement (Cohen’s κ 0.345) between the interpretation based on the two guidelines.

**Table 3.** Inter-rater agreement between original guideline and revised guideline based on the fecal egg count reduction test analyzed using eggCounts. For the purpose of this comparison, the categories normal and susceptible as well as the categories suspected resistance and inconclusive were considered to be equivalent.

|  | Revised guideline |  |  |
| --- | --- | --- | --- |
| Original guideline | Susceptible | Inconclusive | Resistance |
| Normal | 10 | 0 | 6 |
| Suspected resistance | 0 | 0 | 0 |
| Resistance | 0 | 0 | 3 |
| Cohen's $\kappa$ 0.345 | | | |

### Deep amplicon species sequencing of pre-treatment samples

Sufficient numbers of larvae were collected from the nine farms only before treatment, meaning no post-treatment species data were available. The most prevalent species identified were *O. ostertagi* and *C. oncophora*, present on nine and eight farms, respectively (Table S2; Fig. 5). *Cooperia punctata* was found on five farms, *Oesophagostomum radiatum*, *Trichostrongylus axei* and *Oesophagostomum venulosum* on four farms each, *Bunostomum phlebotomum* on three farms, *H. contortus* and *Ostertagia leptospicularis* on two farms and *Cooperia ovina* and *Trichostrongylus colubriformis* on one farm.

**Fig. 5.**
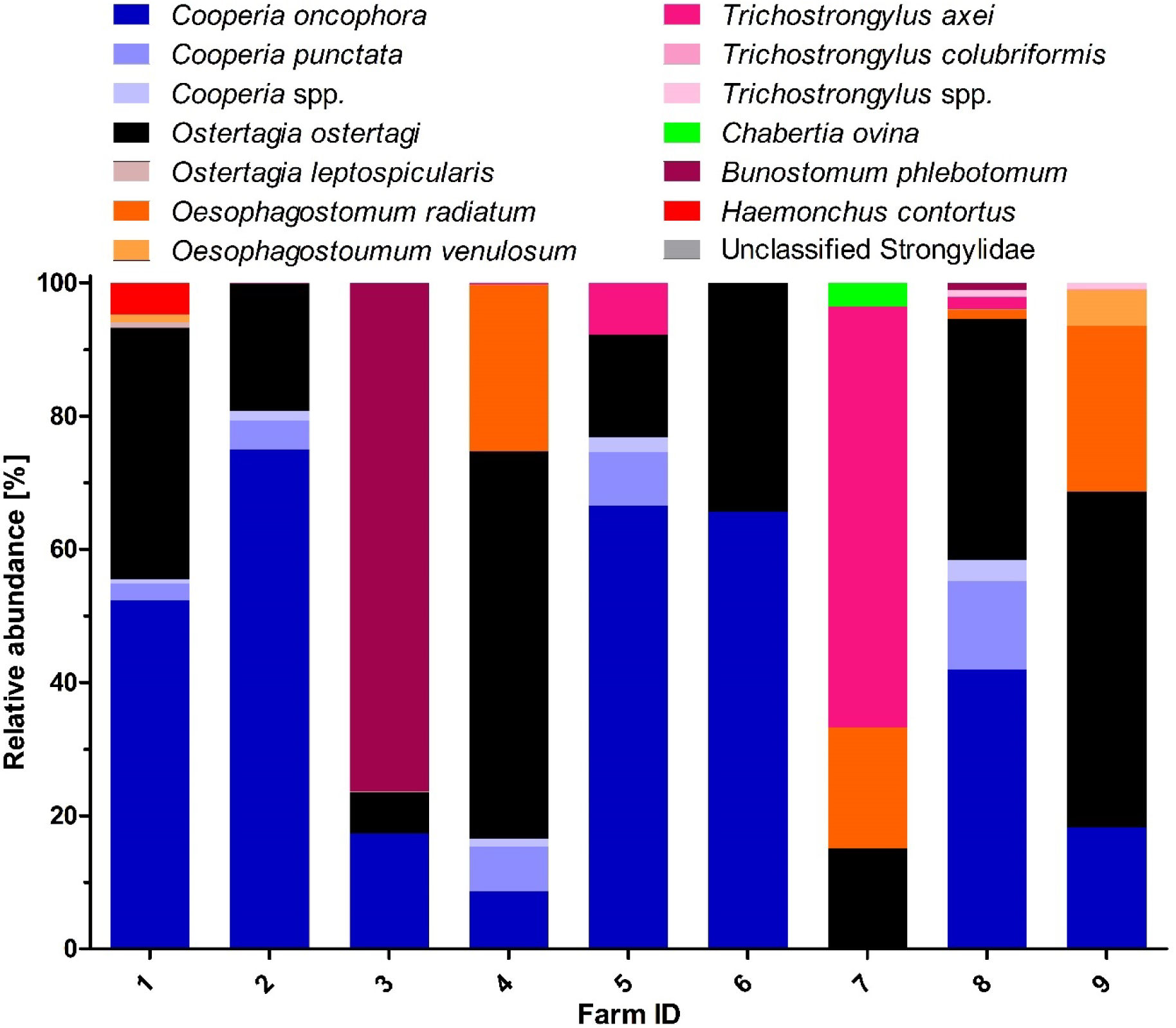
Relative abundance of strongyle species detected in pooled samples from German cattle farms before treatment. A few amplicon sequence variants could not be identified to the species level and were assigned to unclassified *Cooperia* (*Cooperia* spp.), unclassified *Trichostrongylus* (*Trichostrongylus* spp.) or unclassified Strongylidae.

In terms of relative abundance, *C. oncophora* was the most abundant species on the majority of the farms (five out of nine). The most prevalent parasite *O. ostertagi* was most abundant only on two farms. *Bunostomum phlebotomum* and *T. axei* each showed the highest abundance on one farm (Fig. 5). The number of true species detected on the farms (excluding unclassified ASVs) ranged between two (only the most prevalent and abundant parasites *C. oncophora* and *O. ostertagi* on farm 6) to seven (on farm 8). Farms 2, 3 and 4, which showed the poorest results in the FECRT differed considerably in their pre-treatment species composition. On farms 2, 3 and 4, the species with the highest relative abundance were *C. oncophora* (75.0%), *B. phlebotomum* (76.4%) and *O. ostertagi* (58.1%), respectively. The highest relative abundance of *H. contortus* was observed on farm 1, the only farm that kept sheep in parallel with cattle.

## Discussion

The control of GIN relies predominantly on metaphylactic and therapeutic treatment with anthelmintics^4^ and resistance to all drug classes has been reported. In Central Europe, resistance of cattle nematodes is currently not considered to be widespread and reported resistance levels were generally low to moderate. However, it has been shown that frequency of resistance alleles can increase very rapidly upon selection^42^ and once resistance is present, it might rapidly expand if the selection pressure is high. The data presented here show resistance to MLs but particularly to BZs and in this respect are among the very first in Central Europe^16^. On two farms, actually, resistances to both predominantly used drug classes were detected. In comparison to previous reports, this suggest that AR is spreading in cattle populations and that the problem might increase in importance in the future as suggested by a recent meta-analysis^17^.

The FECRT remains the most important test to evaluate the susceptibility/resistance status of gastro-intestinal helminth communities since it is applicable to all drug classes and does not require substantial experimental know-how and standardization as many *in vitro* test methods^43^. However, the original WAAVP guideline for the FECRT^27^ had not been adapted for more than 30 years before a recent revision by Kaplan, et al.^28^. The revised guideline did not only change the guidelines for study design and reporting but also changed the criteria (cut-offs) for identification of resistant populations. In addition, several different statistical approaches have been suggested to analyze FECRT data and to include the different sources of variation^37, 38, 40^. It is important to understand that the application of different criteria/guidelines and statistical approaches results in different interpretation of the same FECRT data. This complicates comparison of results between studies if they have been interpreted using different guidelines or 90% CrIs were calculated using different statistical approaches. We have, therefore, started to systematically compare effects of the application of the original guideline in comparison to the revised guideline and of the usage of different statistical approaches that at least consider different levels of variation according to technical and biological sources of variation^40, 44, 45^. While eggCounts and bayescount both use Bayesian statistics and Markov Chain Monte Carlo approaches to determine the parameters of the model parameters, eggCounts uses a parametric and bayescount a more robust non-parametric model. As typical for non-parametric models, this results in wider 90% CrIs predicted by bayescount as found here but also in previous studies on sheep and goats^29, 46^. When comparing the inter-rater agreement calculated as Cohen’s κ for analysis based on eggCounts and bayescounts, previous studies have reported κ values of 1 and 0.774, which is considerably higher than the 0.518 obtained in the present study. This can be easily explained by the low level of resistance that was detected here for cattle strongyles. This leads to 90% CrIs being very close to the cut-offs of 99% and 95% and even small changes in the CrIs can lead to entirely different interpretation of the data as e.g. for farm 5 with EPR. On this farm, the upper CrL for eggCounts was slightly below and for bayescount slightly above the 99% cut-off leading to interpretation of susceptible and beginning resistance, respectively. Similarly, Krücken, et al.^29^ observed complete agreement for FBZ and IVM treatment with high levels of resistance but poorer agreement for MOX, against which only low-level resistance was found. For the comparison of data obtained employing the original guideline with that obtained when using the revised guideline, a Cohen’s κ of only 0.345 was observed, which is again considerably lower than the values reported before of 0.444 and 0.740^29, 46^. In this comparison, it should be emphasized that many strongyle communities that were considered to be resistant according to the revised guideline were still considered to be susceptible when applying the criteria of the original guideline. This makes it difficult or impossible to compare results of FECRT between studies that applied the original and the revised guideline. Most likely, previous studies using the original guideline underestimated the number of farms with a resistant strongyle community.

The nemabiome analysis revealed that *O. ostertagi* and *C. oncophora* were the most prevalent and abundant species. This is in agreement with other studies from Central Europe using coprocultures^13, 14, 19^ in which these species also dominated. Necropsy studies showed a more diverse strongyle community but of course were only rarely conducted^47, 48^. There were several rather unexpected results, such as the fact of *B. phlebotomum* and *T. axei* being the most abundant on one farm each. In addition, the number of species detected per farm ranged from two (*C. oncophora* and *O. ostertagi*) to seven, including three genera with more than one species, suggesting that morphological differentiation of larvae from coprocultures considerably underestimates species diversity in cattle while nemabiome analysis is much closer to the picture obtained using necropsies.

The fact that resistance against both BZs and MLs was found, especially in parallel on the same farm, shows that efficient control strategies need to be designed and information spread to livestock veterinarians and farmers to improve parasite control. There was no obvious difference between these two and the other farms, as increased treatment frequencies or other management practices, that might lead to faster selection for resistance. Currently, resistance levels are low to moderate and there still are options to prevent that cattle farming in Central Europe moves on to a similar situation as in South America, Australia and New Zealand^9, 10, 11^, where high level multi-drug resistance is widely present. Potential strategies could be e.g. (i) implementation of targeted selective treatment^49^ to leave a large proportion of the parasite populations unexposed to drugs and thus, reduce selection pressure; (ii) introduction of drug combinations to decelerate selection of resistant parasites since it might be difficult to evolve resistance to two drugs simultaneously^50^; (iii) preventing the spread of resistance by effective quarantine and management procedures. It is important to at least now start discussions about preventions measures not only in academia but also with stakeholders. It is important not to miss the point in time when we can still direct the anthelmintic resistance evolution scenario into another direction.

### Conclusions

The present study underscores the emergence of AR to MLs and BZs in cattle nematodes in Germany. Insights in pre-treatment species data yielded a surprising diversity that could provide a more refined understanding of species resistance dynamics with continued observation. However, the inter-farm variability throughout statistical analysis and guidelines results in a more difficult assessment of resistance-data and their interpretation throughout the years. Updated FECRT guidelines, though essential for contemporary assessment, complicate result comparability and emphasize the need for methodological consistency in future AR studies. Immediate adoption of preventive measures, like targeted selective treatment and use of drug combinations, is crucial to mitigate resistance spread. This proactive approach, combined with farmer and veterinary education, can help delay high-level AR development, safeguarding cattle health and productivity in the region.

## Acknowledgements

We would like to thank the participating farmers for their willingness to participate and support the study.

## Data availability

Additional information, data, code for analysis or materials relevant to the research are available upon request to the corresponding author, Georg von Samson-Himmelstjerna at.

## Contributions

PE: Conceptualization, Data curation, Formal analysis, Investigation, Methodology, Validation, Visualization, Writing – original draft, Writing – review & editing. JK: Conceptualization, Data curation, Formal analysis, Funding acquisition, Methodology, Project administration, Supervision, Validation, Visualization, Writing – original draft, Writing – review & editing. SF: Data curation, Formal analysis, Investigation, Validation, Visualization, Writing – review & editing. FH: Data curation, Formal analysis, Validation, Writing – review & editing. CSH: Conceptualization, Writing – review & editing. AN: Conceptualization, Writing – review & editing. WW: Conceptualization, Funding acquisition, Writing – review & editing. WT: Conceptualization, Writing – review & editing. SS: Conceptualization, Funding acquisition, Project administration, Writing – review & editing. RD: Conceptualization, Funding acquisition, Project administration, Writing – review & editing. GvSH: Conceptualization, Funding acquisition, Project administration, Supervision, Writing – original draft, Writing – review & editing.

