## Supplemental material for "Anthelmintic resistance against benzimidazoles and macrocyclic lactones in strongyle populations on cattle farms in northern Germany"

SUPPLEMENTARY MATERIAL

**Table S1** Primers for amplification of internal transcribed spacer for Illumina sequencing

| **Primer names** | **Primer sequences (5' -> 3')** |
|---|---|
| Forward primers | |
| NC1_with_Illumina_Adapter_(0N) | TCGTCGGCAGCGTCAGATGTGTATAAGAGACAGACGTCTGGTTCAGGGTTGTT |
| NC1_with_Illumina_Adapter_(1N) | TCGTCGGCAGCGTCAGATGTGTATAAGAGACAGNACGTCTGGTTCAGGGTTGTT |
| NC1_with_Illumina_Adapter_(2N) | TCGTCGGCAGCGTCAGATGTGTATAAGAGACAGNNACGTCTGGTTCAGGGTTGTT |
| NC1_with_Illumina_Adapter_(3N) | TCGTCGGCAGCGTCAGATGTGTATAAGAGACAGNNNACGTCTGGTTCAGGGTTGTT |
| Reverse primers | |
| NC2_with_Illumina_Adapter_(0N) | GTCTCGTGGGCTCGGAGATGTGTATAAGAGACAGTTAGTTTCTTTTCCTCCGCT |
| NC2_with_Illumina_Adapter_(1N) | GTCTCGTGGGCTCGGAGATGTGTATAAGAGACAGNTTAGTTTCTTTTCCTCCGCT |
| NC2_with_Illumina_Adapter_(2N) | GTCTCGTGGGCTCGGAGATGTGTATAAGAGACAGNNTTAGTTTCTTTTCCTCCGCT |
| NC2_with_Illumina_Adapter_(3N) | GTCTCGTGGGCTCGGAGATGTGTATAAGAGACAGNNNTTAGTTTCTTTTCCTCCGCT |

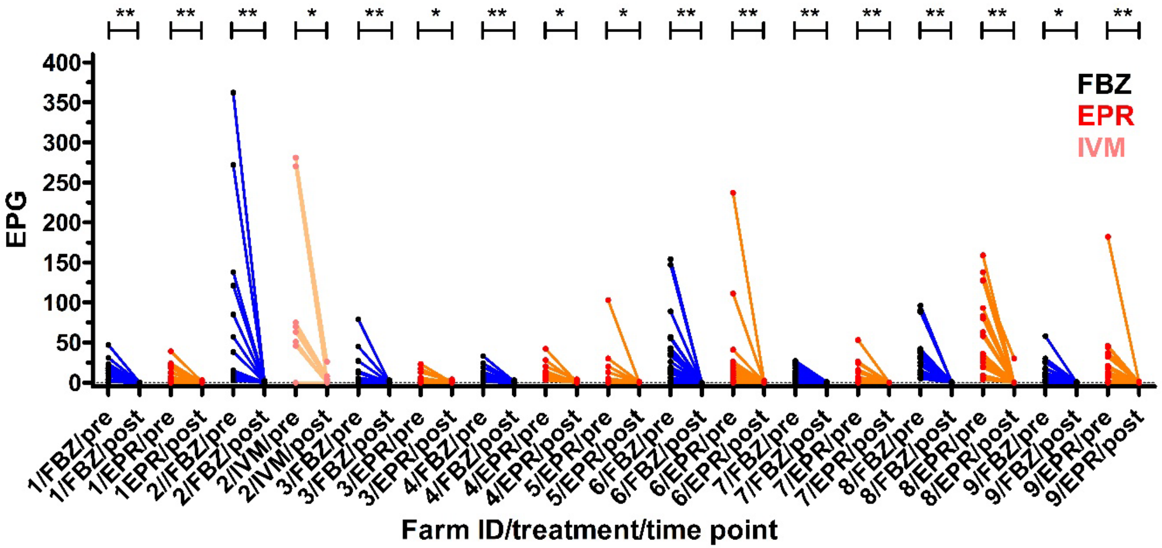


**Fig S1.** Fecal egg counts in eggs per gram feces (EPG) before and after treatment. Pre-treatment samples were collected from the animals on the day of treatment with either fenbendazole (FBZ), eprinomectin (EPR) or ivermectin (IVM). Egg counts of animals positive for strongyle eggs before treatment and the same animals 14 days post treatment were compared with the Wilcoxon matched-pairs signed rank test. **, $p < 0.01$; *, $p < 0.05$. All p values in Fig. S1 and Fig S2 were corrected together for multiple testing using the Holm-Bonferroni method.

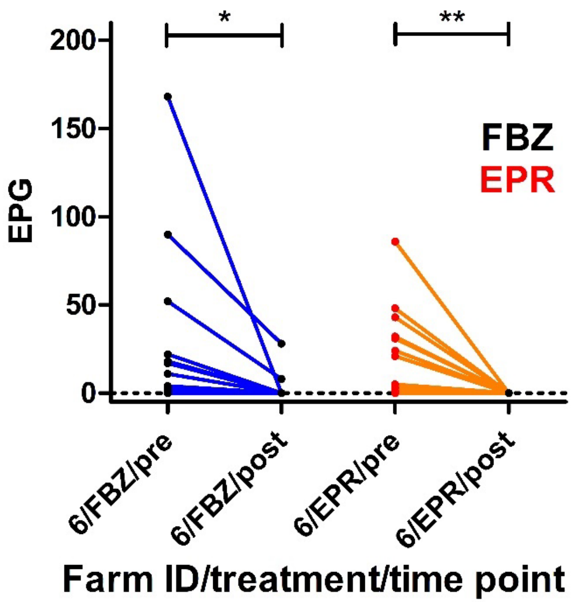


**Fig. S2.** Fecal egg counts in *Nematodirus* spp. eggs per gram feces (EPG) before and after treatment. Pre-treatment samples were collected from the animals on the day of treatment with either fenbendazole (FBZ) or eprinomectin (EPR). Egg counts of animals positive for *Nematodirus spp*. eggs before treatment and the same animals 14 days post treatment were compared with the Wilcoxon matched-pairs signed rank test. **, $p < 0.01$; *, $p < 0.05$. All p values in Fig. S1 and Fig S2 were corrected together for multiple testing using the Holm-Bonferroni method.

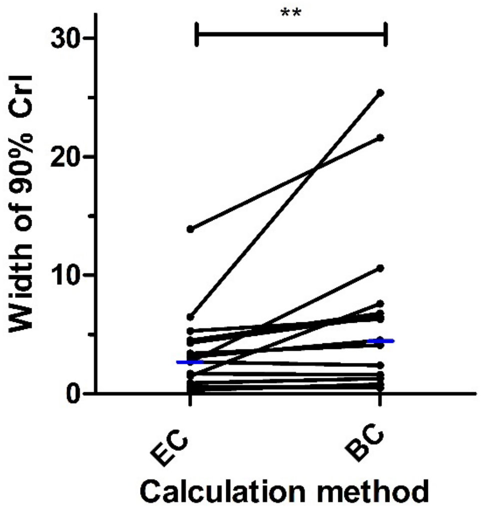


**Fig. S3.** Comparison of the width of the 90% credible intervals (CrIs) between eggCounts (EC) and bayescounts (BC) statistical approaches. Results for the same dataset are connected by lines. Blue lines indicate the medians. Data were compared using a Wilcoxon matched-pairs rank signed test. **, $p < 0.01$.

| Species | N positive | Prevalence (%) | 95% CI (%) |
|---|---|---|---|
| *Cooperia oncophora* | 8 | 88.9 | 56.5 – 98.0 |
| *Cooperia punctata* | 5 | 55.6 | 26.7 – 81.1 |
| *Cooperia* spp. | 5 | 55.6 | 26.7 – 81.1 |
| *Ostertagia ostertagi* | 9 | 100 | 70.1 - 100 |
| *Ostertagia leptospicularis* | 2 | 22.2 | 6.3 – 54.7 |
| *Oesophagostomum radiatum* | 4 | 44.4 | 18.9 – 73.3 |
| *Oesophagostomum venulosum* | 4 | 44.4 | 18.9 – 73.3 |
| *Trichostrongylus axei* | 4 | 44.4 | 18.9 – 73.3 |
| *Trichostrongylus colubriformis* | 1 | 11.1 | 2.0 – 43.5 |
| *Trichostrongylus* spp. | 2 | 22.2 | 6.3 – 54.7 |
| *Chabertia ovina* | 1 | 11.1 | 2.0 – 43.5 |
| *Bunostomum phlebotomum* | 3 | 33.3 | 12.1 – 64.6 |
| *Haemonchus contortus* | 1 | 11.1 | 2.0 – 43.5 |
| Strongylidae | 1 | 11.1 | 2.0 – 43.5 |

**Table S2** Farm prevalence with 95% confidence intervals (CI) of strongyle species identified by deep amplicon sequences in nine German cattle farms before treatment.
